# The P25 protein of *Potato virus X* suppresses accumulation of 24nt siRNAs in the *N. benthamiana* transgenic line 16c

**DOI:** 10.1101/023895

**Authors:** Shahinez Garcia, Christophe Himber, Olivier Voinnet

## Abstract

By transiently expressing viral suppressors of RNA silencing (VSRs) in combination with GFP silencing-inducing constructs in leaves of *Nicotiana benthamiana* line 16c, Hamilton et al. (2002) could establish a positive correlation between the production of 24nt GFP small interfering (si)RNAs in infiltrated leaves and the systemic onset of GFP silencing in remote tissues. In the context of GFP silencing inducers based on replicating *Potato virus X (PVX)*, the P25 protein of PVX was found to specifically inhibit the local accumulation of 24nt GFP siRNAs. In the original paper, there were background pixel pattern duplications in the figure reporting the P25 experiments. We have now repeated these experiments with the original clones and the results presented here confirm those reported in the original paper.

## Introduction

Small interfering RNAs (siRNAs) are the sequence-specific mediators of RNA silencing, a highly conserved gene regulatory mechanism that also serves antiviral functions in several organisms including plants (Hamilton and Baucombe, 1999). To counteract the host gene silencing response, many viruses produce viral suppressors of RNA silencing (VSRs) that target distinct stages of the process (Ding and Voinnet, 2007).

The genome of N.benthamiana line 16c contains a highly expressed GFP transgene that becomes silenced upon transient expression, in single leaves, of exogenous constructs showing sequence homology to the stably expressed GFP transgene (Voinnet and Baulcombe, 1997). Silencing activated in single leaves triggers, in turn, the onset of a systemic signal that silences GFP expression in remote, untreated parts of the 16c plants. Hamilton et al. (2002) showed that both the local and systemic silencing are accompanied by the accumulation of sense and antisense GFP siRNA species, which, when resolved on polyacrylamide gel, migrate as distinct species of ∼21-nt and ∼24-nt, respectively. Remarkably, when co-expressed with silencing-inducing constructs, several VSRs were found to alter the pattern of local siRNA accumulation and the subsequent onset of systemic silencing. In particular, some VSRs suppressed the 24nt siRNA, but not the 21nt siRNA, and in these instances, local silencing was activated but systemic silencing was suppressed (Hamilton et al., 2002; Figure 2-3). These results suggested a positive correlation between the local production of 24nt siRNAs and the onset of a systemic silencing signal in line 16c.

## Results

In the Potato virus X (PVX) genome, the P25 open reading frame (ORF) is part of the so called ‘triple gene block” (TGB), which, together with the coat protein (CP) ORF, is required for viral cell-to-cell and systemic movement; the TGB and CP are both dispensable for virus replication in single cells. A transgene based on replicating PVX engineered to express the GFP ORF and lacking the CP ORF (referred to as PVX-GFP-ΔCP) was shown to trigger local silencing in line 16C; however, the construct was unable to activate systemic silencing. By contrast, when the TGB was removed from PVX-GFP-ΔCP, the resulting replicating virus (referred to as PVX-GFP-ΔTGB-ΔCP) triggered both local and systemic silencing. Ectopic expression of the P25 ORF together with PVX-GFP-ΔTGB-ΔCP was sufficient to block systemic silencing without altering local silencing (Hamilton et al., 2002, Voinnet et al. 2000).

Because the experiments in Figure 2-3 of Hamilton et al. (2002) established a possible correlation between the local production of 24nt GFP siRNAs and systemic silencing in line 16c, the effect of the P25 protein on the accumulation of 24nt siRNA was tested in the context of the viral silencing inducers PVX-GFP-ΔCP and PVX-GFP-ΔTGB-ΔCP, mentioned above. Experiments reported in Figure 4A-B of Hamilton et al. (2002) showed that replication of PVX-GFP-ΔTGB-ΔCP, which does not produce P25, leads to production of both 21nt and 24nt GFP siRNAs, whereas replication of PVX-GFP-ΔCP, which produces P25, promotes accumulation of 21nt siRNAs only. Furthermore, ectopic expression of P25 in combination with PVX-GFP-ΔTGB-ΔCP suppressed accumulation of the 24nt GFP siRNAs without altering the accumulation of the 21nt siRNA. These results therefore reinforced the positive correlation between the local production of 24nt siRNAs and the onset of a systemic silencing signal in line 16c.

However, it was brought to our attention that the original image in Figure 4B of Hamilton et al. (2002) shows background pixel pattern duplications; the duplications did not include any regions of siRNA signal, nor did they obscure any other genuine RNA signal. Nonetheless, to dispel any doubts on the published results, the original PVX-GFP-based constructs and the P25 clone used in Hamilton et al. (2002) were retrieved and a new experiment was conducted in leaves of line 16c. As a control for the production of both 21nt and 24nt GFP siRNAs, we used a sense GFP transgene, a strong inducer of local and systemic silencing (Hamilton et al. 2002). Co-expression of the GFP transgene together with a second transgene encoding the P1 protein of Rice yellow mottle virus provided a control for the specific loss of the 24nt GFP siRNA species, as originally described in Figure 2 of Hamilton et al. (2002). As seen in Figure 1A-B, both the viral (from PVX-GFP-ΔCP) and ectopic expression (together with PVX-GFP-ΔTGB-ΔCP) of P25 was sufficient to suppress the accumulation of 24nt GFP siRNAs without altering that of the 21nt siRNA species, thereby confirming the original results depicted in Figure 4 of Hamilton et al. (2002).

**Figure 1.**
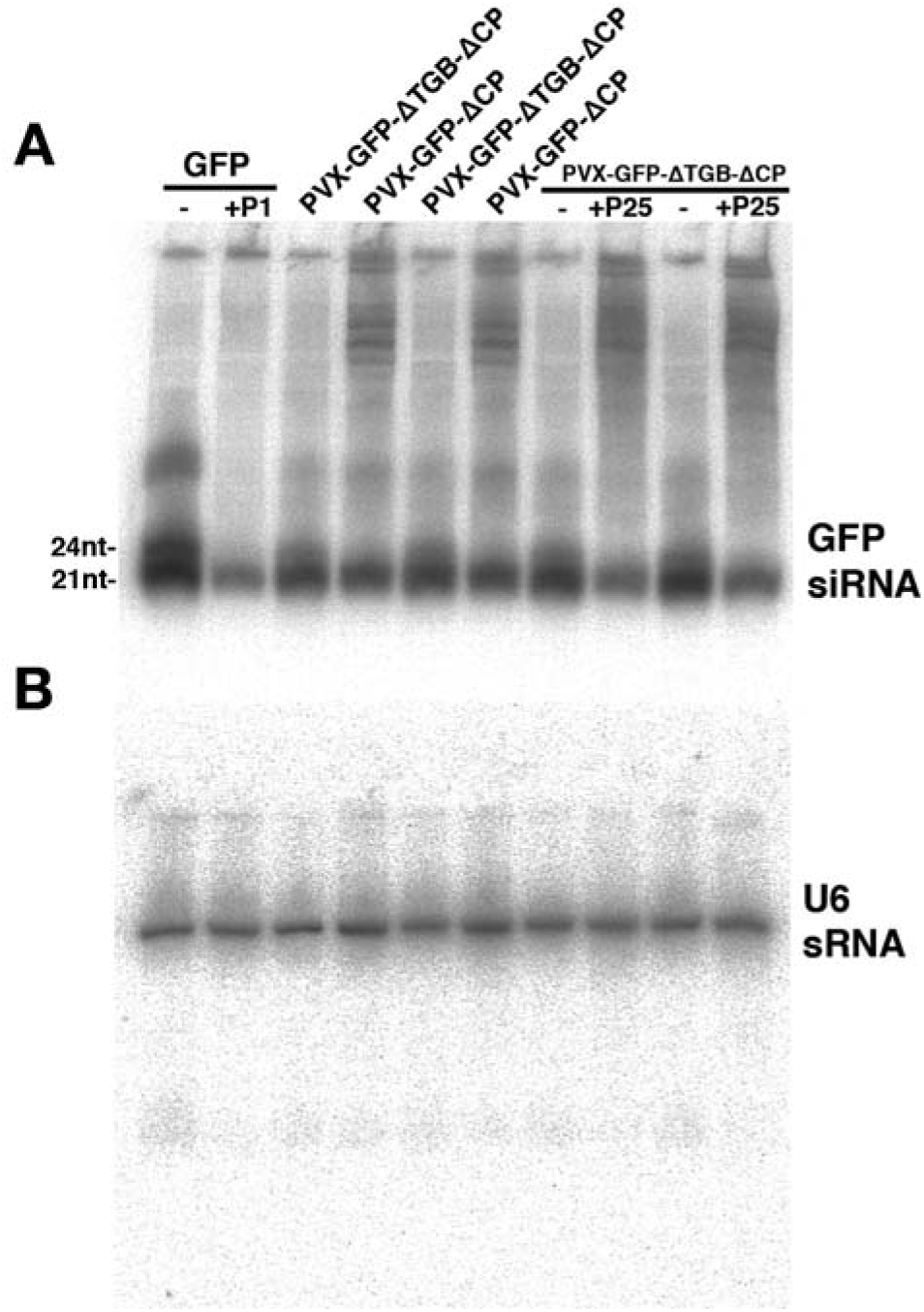
**(A) Low molecular weight RNA analysis.** Total RNA was extracted 5 days after Agrobacterium-mediated transient expression, in leaves of N.benthamiana line 16c, of the indicated PVX-GFP based constructs. 15μg of total RNA were separated on acrylamide gel (mini-protean casting system), blotted and hybridized with a random-primed DNA probe corresponding to the 5’ part of the GFP open reading frame. RNA extracted from a leaf infiltrated 5 days earlier with the GFP construct was used as a positive control for the accumulation of both 21nt and 24nt GFP siRNA species. RNA extracted from a leaf co-infiltrated with GFP and the P1 suppressor of silencing was used as a positive control for the specific loss of the 24nt GFP siRNA species. Note the specific effect of virally or ectopically expressed P25 on the 24nt viral-derived GFP siRNA species. **(B)** The membrane in (A) was stripped and re-hybridized with an oligonucleotide probe specific for the constitutive U6 small RNA, providing an indirect assessment of the quantities of small RNAs for each sample tested.

## Material and methods

Constructs PVX-GFP-ΔCP, PVX-GFP-ΔTGB-ΔCP, P25, P1, GFP, and N. benthamiana line 16c were as described in Hamilton et al. (2002). Agrobacterium-mediated transient expression of the various constructs in individual leaves of the N.benthamiana 16c line was performed as described in Hamilton et al., 2002; 5 days post-infiltration, total RNA was extracted from the infiltrated leaves using TRIZOL (Invitrogen) and 15μg were separated on acrylamide gel (BIORAD mini-protean casting system), blotted, chemically cross-linked (Pall and Hamilton, 2008) and hybridized with a random-primed DNA probe corresponding to the 5’ part of the GFP open reading frame. The membrane was exposed for 2h under phosphorimager.

